# Transcriptional bursting, gene activation, and roles of SAGA and Mediator Tail measured using nucleotide recoding single cell RNA-seq

**DOI:** 10.1101/2024.03.08.584165

**Authors:** Jeremy A Schofield, Steven Hahn

## Abstract

A time resolved nascent single-cell RNA-seq approach was developed to dissect gene-specific transcriptional bursting and the roles of SAGA and Mediator Tail (the activator-binding module). Most yeast genes show near-constitutive behavior while only a subset of genes show high mRNA variance suggestive of transcription bursting. Bursting behavior is highest in the coactivator redundant (CR) gene class (dependent on both SAGA and TFIID) and is strongest in TATA-containing CR genes. Applying this approach to analyze gene activation, we found that basal histone gene transcription is in a low level, low-noise constitutive mode while the activated state unexpectedly shows an increase in both the fraction of active promoters and a switch to a noisy and bursty transcription mode. Rapid depletion of either SAGA or Mediator Tail suggests that both factors play an important role in stimulating the fraction of active promoters at CR genes, with a variable gene-specific role in transcriptional bursting.

## Introduction

Eukaryotic protein-coding genes can be transcribed either constitutively, characterized by stochastic uncorrelated initiation events, or in bursts of transcription initiation separated by stochastic on/off periods^1–3^. Bursting is described by several kinetic parameters that vary by gene and chromatin environment. These parameters include the frequency of transition to the active promoter state, the duration of active periods, and the number of initiation events during the active period (burst size)^4^. The frequency of activation is generally thought to be controlled through enhancer activity in metazoans^5^, while the size and variance of bursting has been attributed to core promoter elements. Transcription factors and chromatin regulators have also been demonstrated to regulate the timing and magnitude of transcriptional bursts in yeast^6,7^. Genes differ considerably in their bursting behavior^8,9^ and, although “burstiness” is not easily described by a single kinetic term, bursty gene expression is characterized by large quantities of RNA produced in a single cell over a short time leading to high mRNA variance per cell, referred to as intrinsic transcriptional noise^10,11^. This transcriptional heterogeneity is most prominent for genes in stress response pathways and in cancer cells^12,13^. The question of how bursting relates to gene regulation has been explored in metazoans where studies found that modulation of promoter activation frequency is the dominant form of regulation^14^. However, subsets of genes can be regulated through modulation of burst size^15,16^. For small numbers of genes, studies of gene-specific transcription kinetics using live cell microscopy have illuminated roles for specific regulators and DNA sequence contexts associated with variance in bursting^3,17^, but the generality of bursting behavior and gene activation mechanisms for most genes remains an open question.

In most systems studied, TATA-containing promoters are often associated with high variance, noise, and bursting ^18–20^. One interpretation of this finding is that TATA promotes retention of a stable “scaffold complex” after Pol II escape (a subset of basal transcription factors bound to the promoter), leading to rapid reinitiation and bursting.^18,21^ However, the finding that TATA-containing genes display higher transcriptional noise may be confounded by the fact that other gene properties overlap with TATA, including high expression^22^ and gene-specific regulation by transcriptional cofactor complexes including SAGA^23,24^ and Mediator Tail (MED)^25,26^.

In yeast, three classes of genes have been defined based on their cofactor dependence. The coactivator redundant gene class (CR genes; ∼13% of protein coding genes)^23,24^ are partially dependent on both SAGA and TFIID and rapid co depletion of these factors leads to a severe defect in transcription of the CR genes. CR genes are enriched for stress response genes and ∼60% of CR gene promoters contain a TATA. ∼44% of the CR genes are also sensitive to rapid depletion of the MED Tail module (the activator-binding domain of yeast MED) and ∼70% of these Tail-dependent gene promoters contain TATA ^25,26^. Conversely, the remaining 87% of genes (TFIID-dependent genes) are enriched for housekeeping genes, only ∼20% of these promoters contain TATA, and transcription decreases upon rapid TFIID (but not SAGA) depletion^23^. Finally, Transcriptional cofactors display specificity for UAS sequences in yeast^27^ and for enhancers in metazoans^28,29^ and, despite the observed connection between enhancers and transcriptional activation frequency, the roles of transcriptional cofactors in gene-specific transcriptional bursting behavior have not been studied systematically. Current limitations in studying gene-specific bursting for large numbers of genes include low throughput in microscopy approaches and the lack of time resolution in standard scRNA-seq data. However, recent advances in RNA metabolic labeling methodology^30,31^ and single-cell RNA-seq^32,33^ now permit time-resolved estimates of transcription in single cells.

To address transcriptional kinetic parameters and how these are affected by cofactors on a genome-wide scale, we use a time-resolved single cell RNA-seq (nucleotide recoding-scRNA-seq, NR-scRNA-seq) approach to measure transcription over short time periods across the yeast transcriptome. Nucleotide recoding approaches allow for the simultaneous measurement of nascent and pre-existing RNAs, which can be leveraged to estimate steady-state transcript half-lives, as well as capture the number of newly made RNAs over time^34,35^. Using this approach, we found bursting behavior associated with specific coactivator regulatory class genes as well as specific promoter elements. We also examined how kinetic parameters change upon gene activation and how rapid depletion of SAGA and MED Tail alter transcription properties genome-wide. Our approach allows rapid and large-scale measurement of transcription properties and should be adaptable to many cell types.

## Results

### Genome-wide measurement of transcription noise and F_on_ using NR-scRNA-seq

To measure kinetic parameters for the set of expressed protein-coding genes, we adapted scRNA-seq to include metabolic RNA labeling and nucleotide recoding chemistry (NR-scRNA-seq) (**Fig 1A**). We performed a time course of 4-thiouracil (4TU) metabolic labeling (5, 10, 15 min) in yeast cells undergoing log growth, treated cells with alkylation chemistry, and prepared droplet-based RNA-seq libraries (10X Genomics). The NR-seq approach modifies 4TU nucleobases in newly transcribed RNAs, resulting in a T-to-C mutational signature in cDNA derived from new RNAs. To determine the efficacy of nucleotide recoding, we first analyzed single cell datasets in bulk. We used a Bayesian modeling approach (bakR^35^) to estimate the T-to-C mutation rate in nascent vs. pre-existing RNAs. We estimate the mutation rate in nascent transcripts for labeled samples ranges from 4.7%-5.7%, while the background mutation rate in pre-existing RNAs is 0.04%. Using a mixed-binomial approach to estimate the fraction of newly transcribed RNAs over the label period^30,35^, we determined the half lives of thousands of polyA transcripts. Transcript half-lives estimated from our three independent timepoints (**Table S1**) are highly reproducible (**Fig S1A**), the median half-life of 8.2 minutes agrees well with prior estimates in yeast and displays the known correlation with codon optimality (**Fig 1B, Fig S1B**). To gain an understanding of the frequency of promoter activation for yeast genes, we next estimated the fraction of cells actively expressing each gene (F_on_) across the timepoints and found a global increase in median F_on_ over time (**Fig 1C**). Due to the low number of new RNA observations for lowly expressed transcripts, our F_on_ measurements are likely underestimates. Despite this, our F_on_ estimates are highly reproducible between independent experimental runs (**Fig S2**), and scale appropriately with metabolic labeling time, validating the consistency in temporal information gained in this approach.

**Figure 1.**
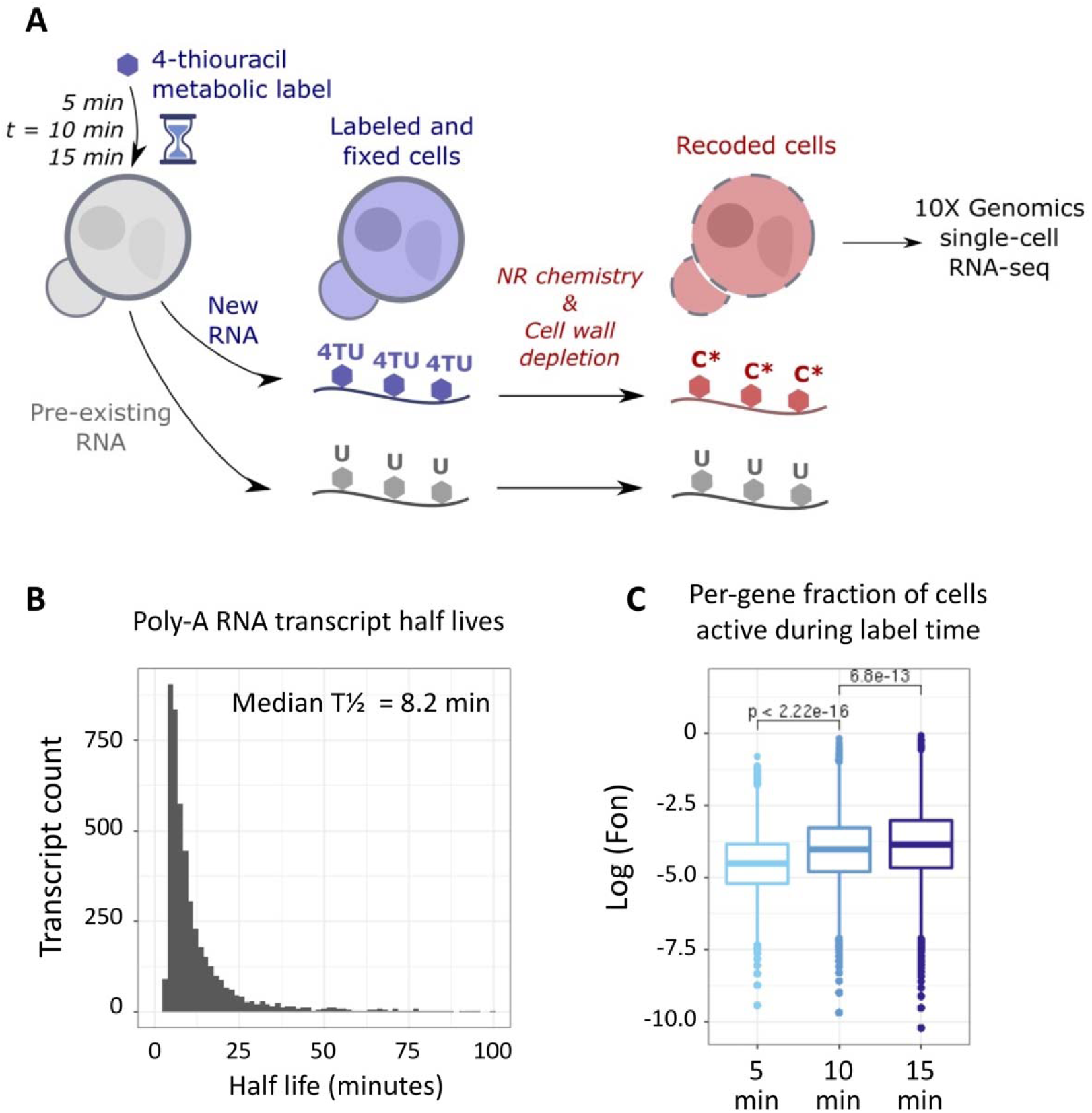
Nucleotide recoding single cell RNA-seq (NR-scRNA-seq) approach to measure transcription in yeast cells. **A**) Scheme of metabolic labeling, cell fixation, NR chemistry and single-cell RNA-seq approach. **B**) Distribution of poly-A transcript half-lives, calculated from bulk fraction-new mRNA estimates derived from the 10 minute labeling timepoint. **C**) Estimates of per-gene fraction on (F_on_) for 5, 10, and 15 minute labeling timepoints. Statistic shown is Wilcoxon rank sum test.

To analyze transcriptional noise strength for thousands of yeast genes, we estimated the number of new RNAs synthesized over time per gene and per cell using a binomial probability approach (see methods, **Fig S3**). We then used these new RNA count estimates to calculate the mean-normalized variance (Fano factor) of new RNAs across many cells. Assuming a Poisson process where transcription initiation events are independent, we expect variance to be equal to the mean (Fano = 1). A Fano factor greater than 1 indicates higher variance than the mean and a non-independent relationship between transcription initiation events. As expected, Fano captures the variance in mRNA abundance observed for different genes as revealed by NR-scRNAseq (**Fig 2A**) and genes with high Fano are associated with transcription bursting^36^. Importantly, Fano estimates agree well across timepoints and experimental strains (**Fig S4A**). Strikingly, most genes display relatively low Fano levels (median Fano = 1.19, n = 2968)), with a minority of genes displaying exceptionally high Fano (Fano > 2, n =16), suggesting a high level of transcriptional bursting (**Fig 2B**). This is consistent with prior results where most yeast genes assayed displayed constitutive behavior (low variance in RNA and protein) with a subset of genes displaying high variance.^3,17,37,38^

**Figure 2.**
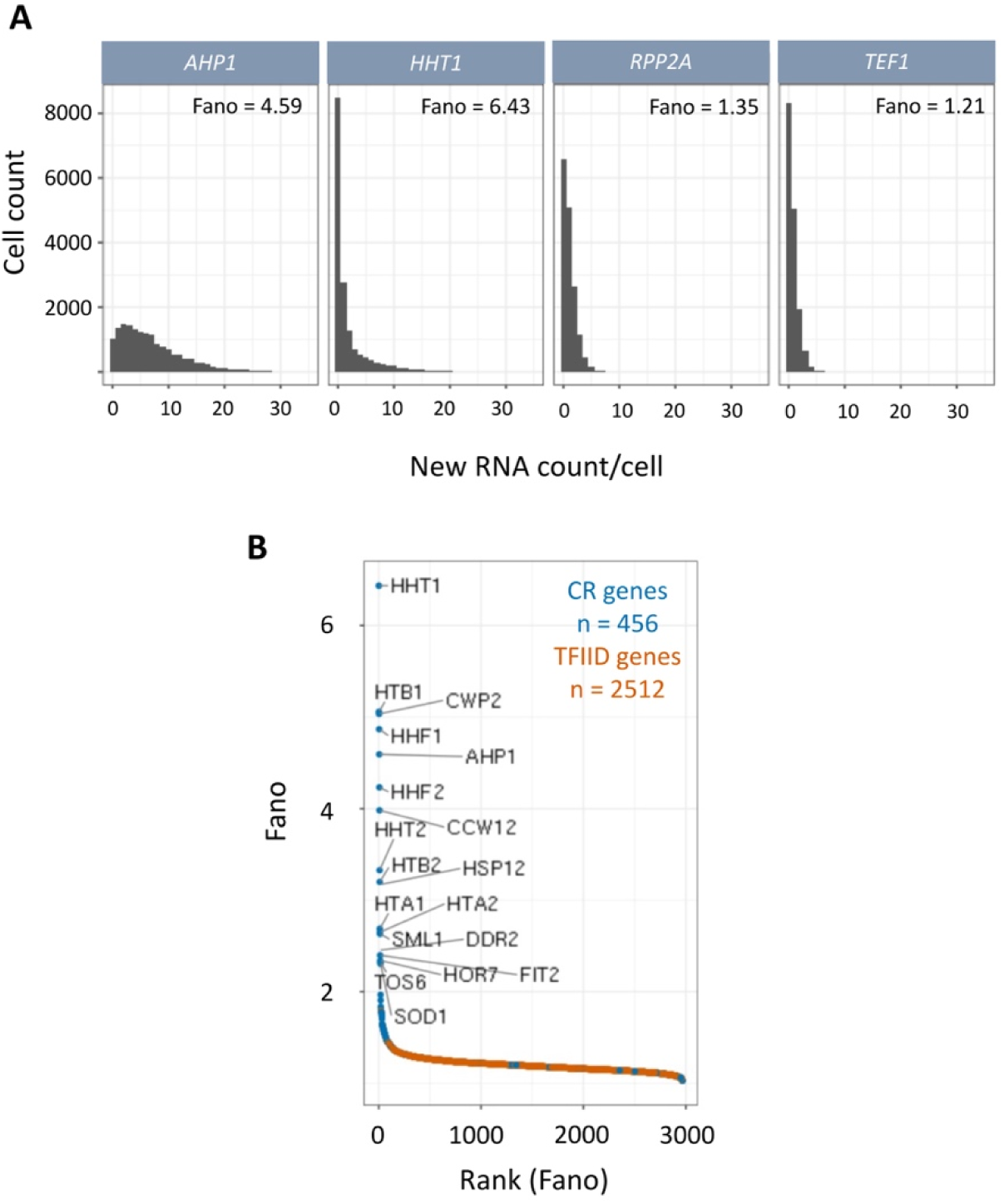
Wide variance in mRNA abundance observed at different genes. **A)** Estimated distribution of new RNA counts per cell for selected high expression genes. Estimated Fano value is displayed for each gene. **B**) Rank order plot of calculated Fano values for approximately 3,000 yeast genes. Genes are colored by coactivator class^23^ with selected high Fano genes labeled.

From our results, bursty genes are highly expressed, although even among highly expressed genes, we observe a wide range of Fano values, suggesting burstiness is not related to expression level alone (**Fig 3A**, **Table S2**). CR genes on average display higher Fano than TFIID genes across 3 of 4 expression quartiles (**Fig 3A**), and we found that CR genes containing TATA boxes generally display higher Fano than CR genes without TATA boxes (**Fig 3B**). However, TATA is not the only important feature as CR genes containing TATA boxes show higher variance than TFIID genes with TATA boxes (**Fig 3B**), suggesting that factors regulating CR transcription also play an important role in promoting bursting behavior. Interestingly, Med Tail-dependent genes have comparable Fano whether or not they contain a consensus TATA (**Fig 3C**), however, nearly all Tail-dependent promoters lacking a consensus TATA contain a closely related TATA derivative^27^. Finally, we observe that promoters with long residence time of the basal transcription factor TFIIF^39^, presumably due to a longer-lived activated state, display on average higher Fano values (**Fig S4B**), providing a connection between our measured transcriptional variance and an independent kinetic analysis of basal factor-promoter stability. Together our results indicate that, on a genome-wide scale, both gene class and promoter elements contribute to transcriptional noise and bursting behavior.

**Figure 3.**
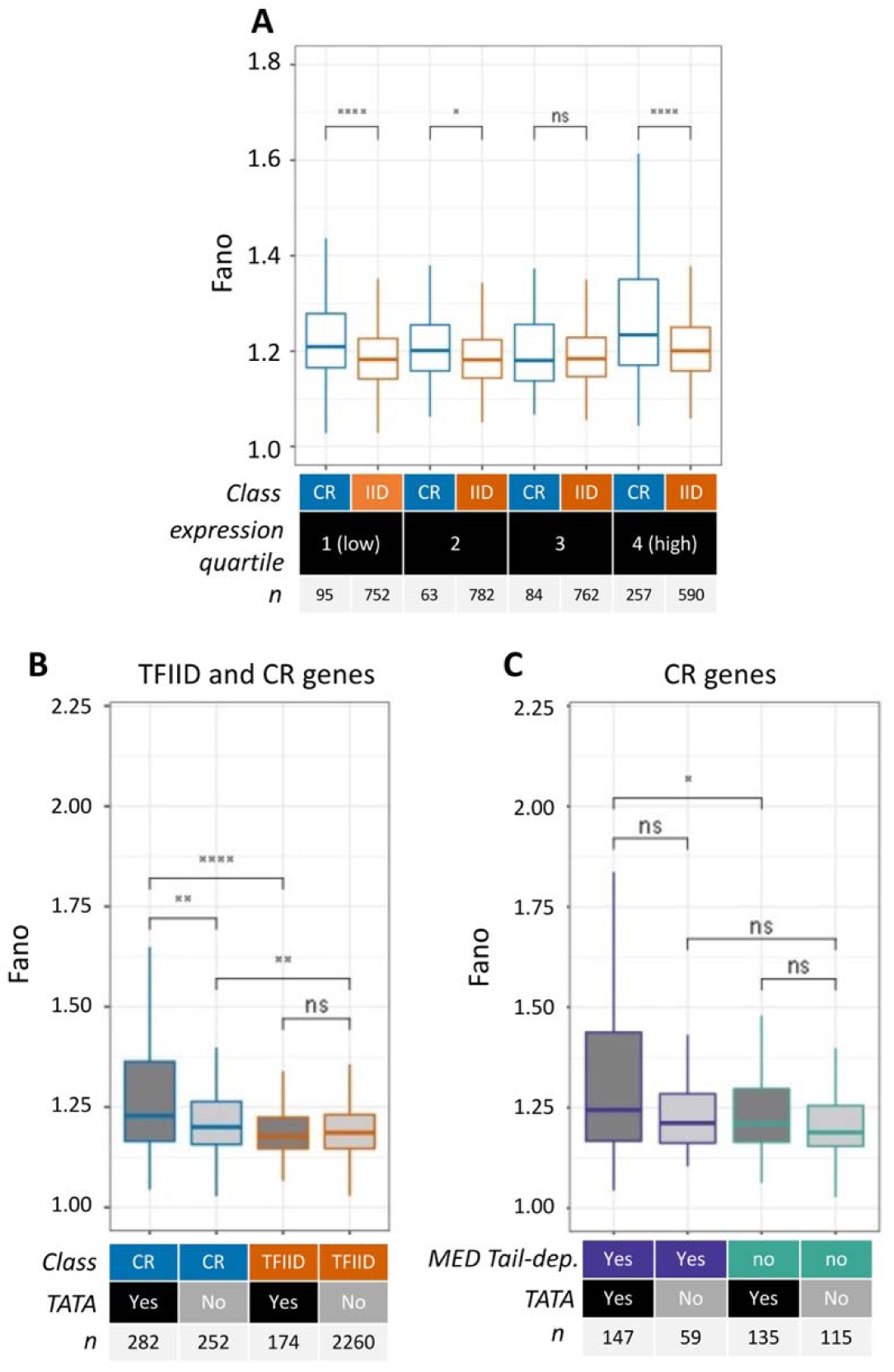
Genomic properties corresponding to Fano values determined by NR-scRNA-seq in yeast. **A**) Boxplot displaying Fano values (outliers not plotted), comparing CR and TFIID genes across expression quartiles. Statistic shown is Wilcoxon rank sum test, number of genes in each category is displayed as “n”. **B**) Boxplot displaying Fano values (outliers not plotted) comparing CR and TFIID genes with and without promoter TATA boxes^23^. Statistic shown is Wilcoxon rank sum test, number of genes in each category is displayed as “n”. **C**) Boxplot displaying Fano values for CR genes (outliers not plotted) comparing MED Tail-dependent and MED Tail-independent genes^26^ with and without promoter TATA boxes^23^. Statistic shown is Wilcoxon rank sum test, number of genes in each category is displayed as “n”. For all figure panels, significance levels are defined as ∗∗∗∗p ≤ 0.0001, ∗∗∗p ≤ 0.001, ∗∗p ≤ 0.01, ∗p ≤ 0.05, ns p > 0.05.

### Activation of cell cycle regulated genes

We next assessed how transcription activation relates to kinetic behavior, and if gene-specific noise is intrinsic to the promoter or varies by the activation state of the gene. As an example of gene activation mechanisms, we examined changes in transcription parameters of cell-cycle specific genes. We first grouped cells from our 10 min 4TU dataset into cell cycle states based on new RNA counts using K-means clustering on a set of 297 curated cell cycle regulated genes^40^ (Methods). Although clustering will not precisely define the cell cycle state, and there is some temporal overlap between cell cycle states due to the 10 min metabolic labeling time, this approach successfully recapitulates known cell cycle dependent transcriptional regulation based on transcript abundance (**Fig 4A**). We found that generally, genes whose expression levels peak in each cell cycle stage also display the highest F_on_ for that corresponding cell cycle stage (**Fig S5A**), consistent with the previously observed connection between gene expression output and the fraction of active promoters. We then compared the F_on_ to Fano for genes at each stage of the cell cycle to assess how mRNA variation changes under differing activation states. Consistent with prior knowledge, we observe an increase in F_on_ for histone genes (**Fig. 4B**) increasing in G1 phase, peaking in S phase, and decreasing in G2 phase. Interestingly, we observe similar high levels of variance (Fano) in G1, S, and G2 cells, compared to the relatively low levels of variance in M and M/G1 cells (**Fig 4B**). Although histone transcription is cell-cycle regulated, we observe ∼15-20% of cells actively transcribing histone genes during the M and M/G1 cell cycle stages, suggesting that histone genes are transcribed but are in a less active state during these latter cell cycle stages. In addition, non-cyclic genes display relatively uniform levels of Fano across the cell cycle (Fig **S5B**), suggesting that the high variance state for histone genes in G1, S, and G2 cells is not influenced by global changes across the cell cycle. Our combined results suggest that the histone genes are transcribed at a low level in a low-noise constitutive mode during M and M/G1 while the activated state is characterized by an increased F_on_ as well as switching to a noisy and bursty transcription mode in G1, S, and G2.

**Figure 4.**
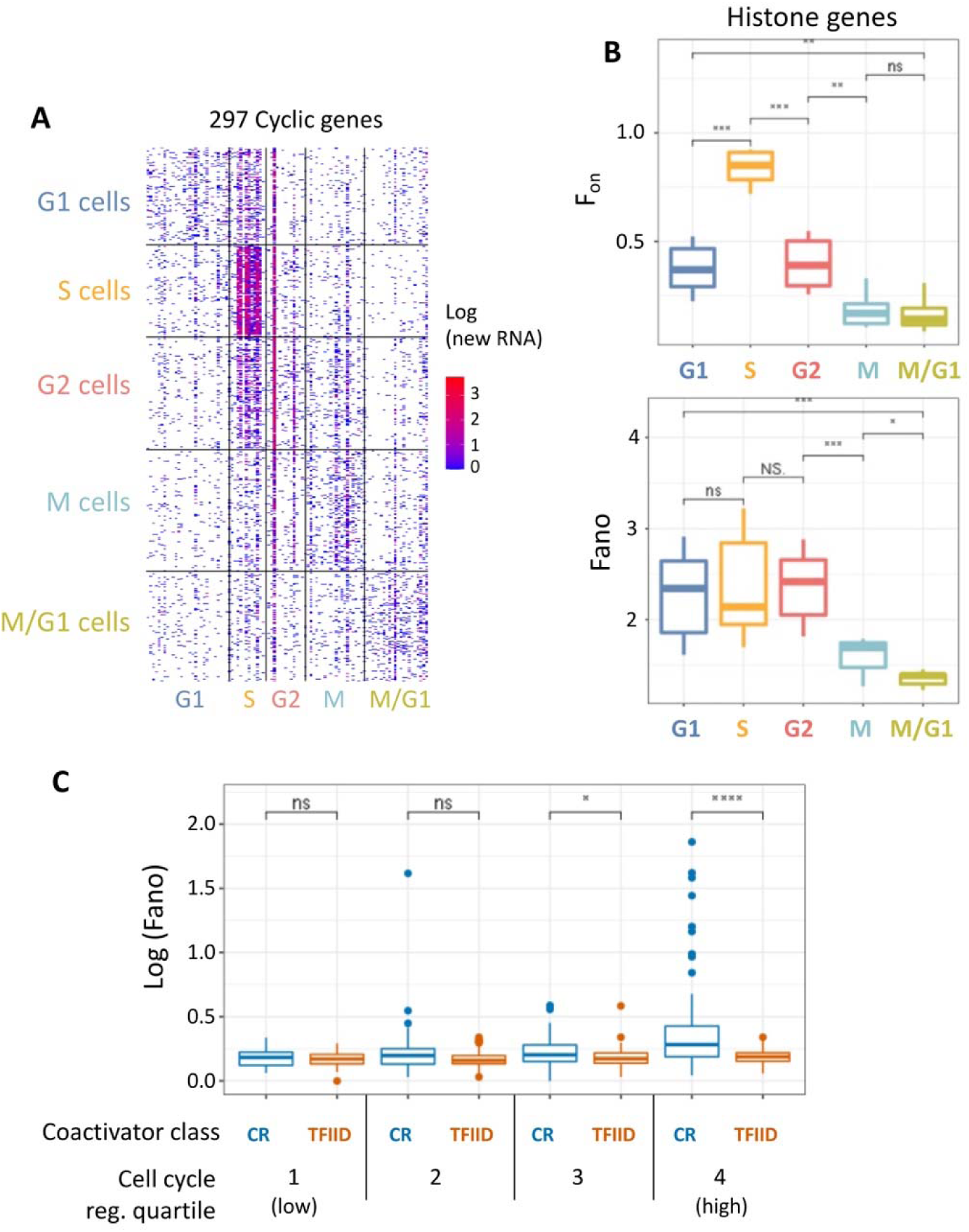
Kinetics of transcription across the yeast cell cycle. **A**) New RNA count matrix for cells assigned to differing cell cycle stages (grouped, y axis), based on k-means clustering of cyclic genes. Genes are grouped on x axis by approximate cell cycle state of peak expression^40^. **B**) F_on_ estimates (top) and Fano estimates (bottom) for histone gene expression in cells assigned to differing cell cycle stages. **c**) Fano estimates for CR vs. TFIID class genes^23^, divided into quartiles based on cell cycle regulation score^40^. Statistic shown is Wilcoxon rank sum test, significance levels are defined as ∗∗∗∗p ≤ 0.0001, ∗∗∗p ≤ 0.001, ∗∗p ≤ 0.01, ∗p ≤ 0.05, ns p > 0.05.

We then assessed whether the degree of cell cycle regulation leads to differences in transcriptional noise. We binned genes according to degree of cell cycle regulation^40^ and found that both CR genes and TFIID genes with low cell cycle regulation (quartile 1) display similarly low Fano (**Fig 4C**). However, for genes with high cell cycle regulation (quartile 4), CR genes display significantly higher Fano than TFIID genes, suggesting that a high variance activated state is a feature of the highly regulated CR gene class.

### Effects of cofactor depletion on transcription parameters

Since we observed differences in transcriptional variance across gene regulatory classes, we interrogated the gene-specific effects on noise and gene activation after rapid depletion of transcription cofactor subunits using the auxin-inducible degron (AID) system^41^. For this analysis, we consider transcriptional bursting to be associated with high mean expression per active cell (MPC) and high transcriptional variance (Fano), and gene activation to be associated with the fraction of cells actively transcribed over the labeling period (F_on_). The role of the SAGA complex was interrogated by co depletion of the core SAGA subunits Spt3 and Spt7 and the role of MED Tail by depletion of the Med15 subunit (**Fig S6A**). Upon Med15 depletion, we see strongest defects in F_on_ for Tail-dependent genes, with modest changes in MPC and Fano for this same gene set **(Fig S6B)**. SAGA depletion results in clear changes in F_on_, MPC and Fano for the larger set of CR genes **(Fig S6C)**.

Examining genome-wide changes following Med15 and Spt3/7 depletion (**Fig 5A-B**), we observe a strong correlation (R^2^ = 0.93, 0.94; linear model fit) between changes in bulk transcription and F_on_, and a modest correlation (R^2^ = 0.34, 0.22) between changes in bulk transcription and MPC. Consistent with our hypothesis of MPC and Fano being indicative of transcriptional bursting, we observe a moderate correlation (R^2^ = 0.57, 0.62) between the two parameters upon Med Tail and SAGA depletion. These trends are consistent when considering only MED-Tail dependent genes following Med15 depletion (**Fig S7A**), as well as CR genes (**Fig S7B**) and TFIID genes (**Fig S7C**) following Spt3/7 depletion. Taken together, our observations support a model of MED-Tail and SAGA function playing an important role in stimulating the fraction of active genes (reflected in F_on_), with a modest role in transcriptional bursting behavior (reflected in MPC and Fano).

**Figure 5.**
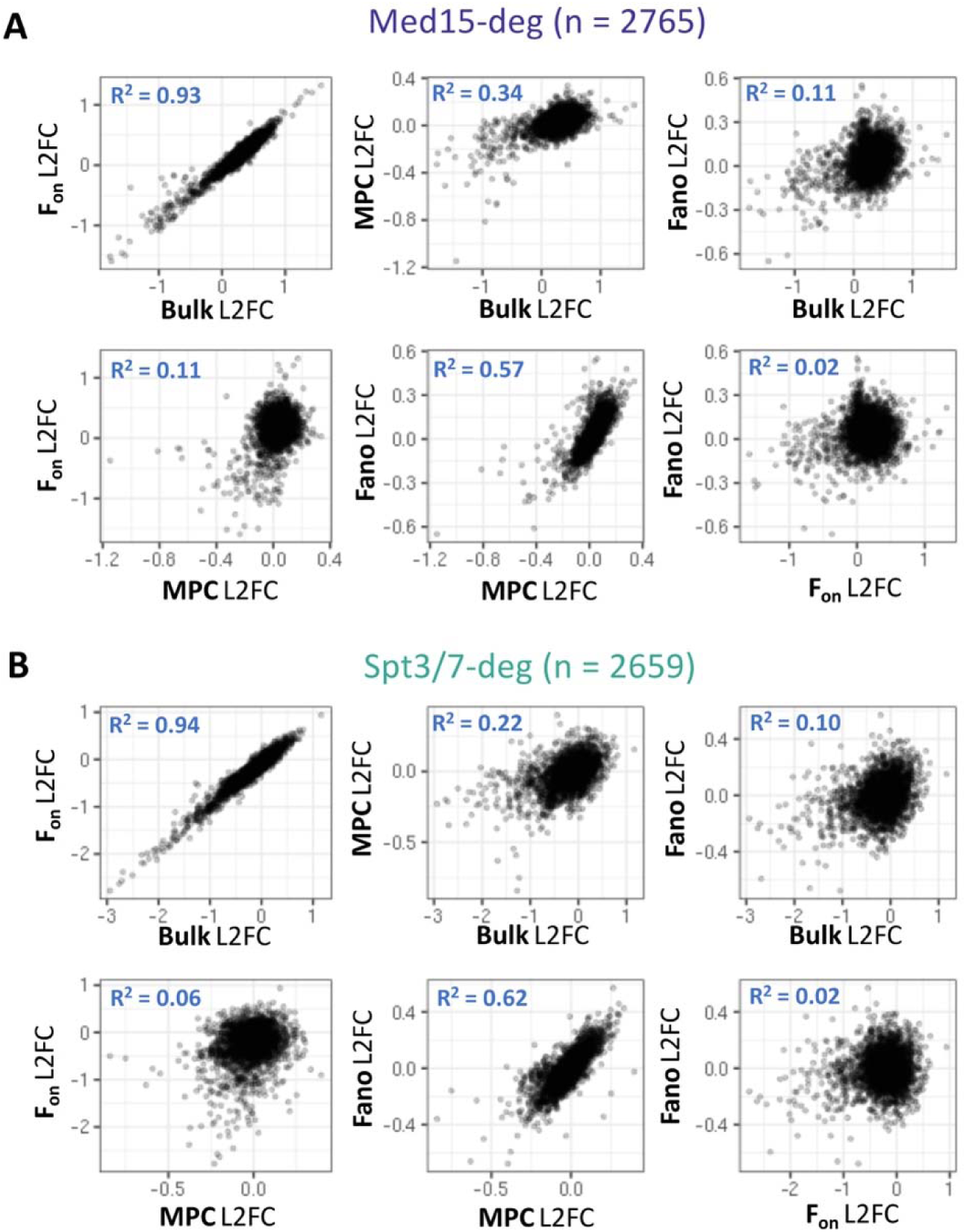
Perturbation of transcriptional kinetic parameters following acute depletion of MED Tail and SAGA coactivator complexes. **A**) Pairwise comparisons between log2-fold changes (L2FC) in bulk transcription, F_on_, MPC and Fano following acute depletion (30 min) of MED Tail (Med15 subunit). R^2^ values shown are adjusted R-squared values derived from linear model fits. **B**) Pairwise comparisons as in A), but measurements after acute depletion of SAGA (Spt3/7 subunits).

In addition to the above analysis of coactivator dependence, it was also informative to examine the behavior of other gene classes and specific genes as we found examples of gene-specific changes in F_on_, MPC and Fano following coactivator depletion. We first focused our analysis on histone genes following Tail or SAGA depletion. Yeast histones genes are chromosomally arranged in pairs as divergent tandem promoters. While all histone genes are in the CR gene class and dependent on SAGA^23^, only two of the four pairs of histone genes are dependent on MED Tail^26^. Following Tail depletion, we observed defects in bulk transcription, F_on_, MPC, and Fano for the Tail-dependent but not the Tail-independent histone genes (**Fig 6A**). Following SAGA depletion, we observe defects in all four parameters for all histone genes, and these defects are largely comparable between Tail-dependent and Tail-independent genes. These results illustrate how transcriptional regulators play distinct roles at different subsets of genes, which can affect transcriptional output and variance.

**Figure 6.**
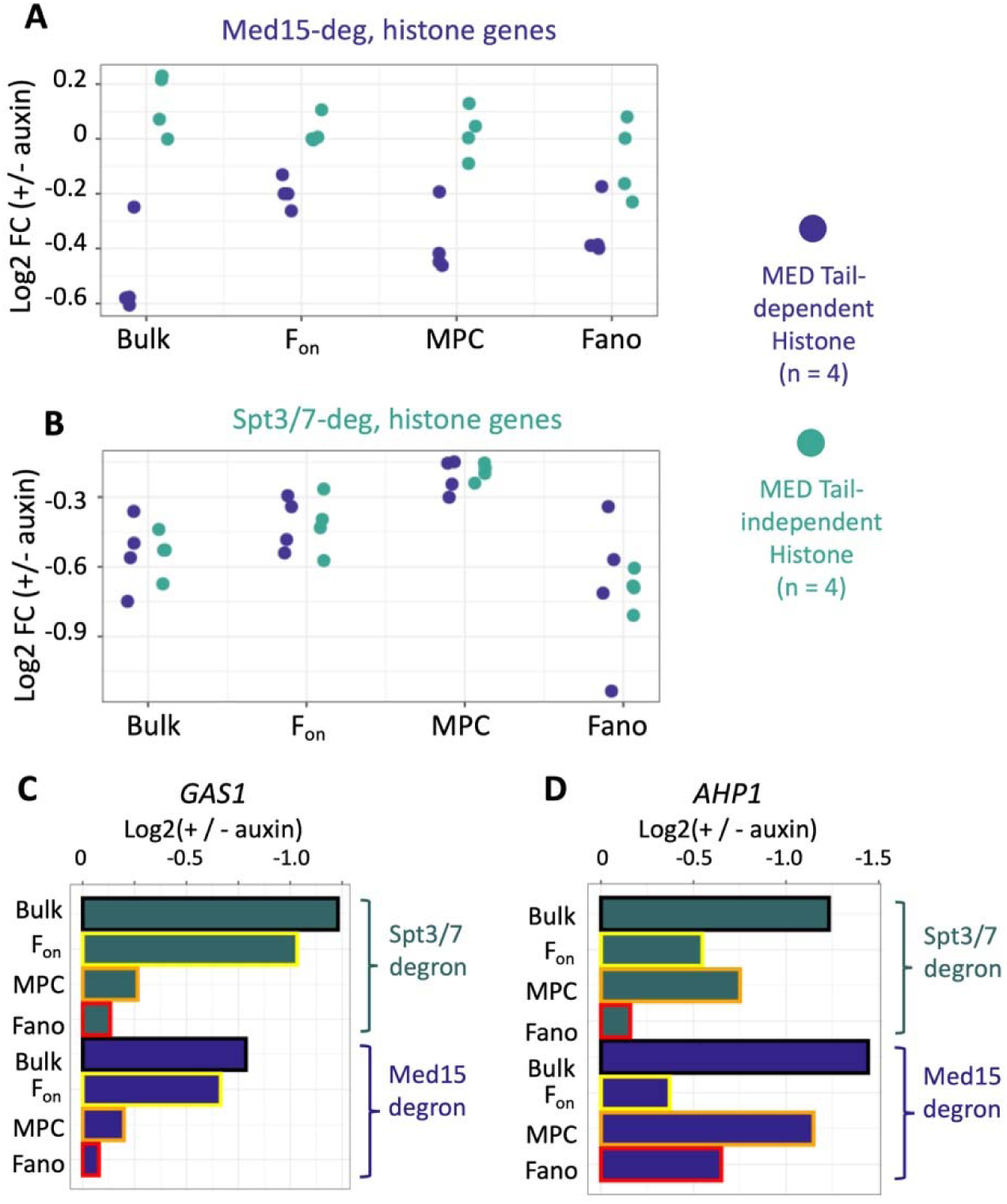
Gene-specific perturbations in transcriptional kinetic parameters following coactivator depletion. **A-B**) Log2-fold change in bulk transcription, F_on_, MPC, and Fano following Med15 depletion (a) or Spt3/7 depletion (b) for the eight histone genes. MED Tail-dependent genes are shown in purple, while MED Tail-independent genes are shown in teal. **C-D**) Log2-fold change in bulk transcription, F_on_, MPC and Fano for (**c**) *GAS1* and (**d**) *AHP1* genes following Spt3/7 or Med15 depletion.

To further illustrate this point, we examined changes in parameters following coactivator depletion for specific non-histone genes. For ribosomal biogenesis (RiBi) genes, SAGA depletion generally results in little effect on bulk transcription and the three measured parameters (**Fig S8A**), while Tail depletion generally results in a slight increase in both bulk transcription and all measured parameters (**Fig S8B**). However, for ribosomal protein (RP) genes, SAGA depletion results in a decrease in transcription, MPC, and F_on_, with a slight decrease in Fano (**Fig S8A**). Conversely, MED Tail depletion results in an increase in bulk transcription and MPC, with a substantial increase in Fano (**Fig S8B**). These results are consistent with prior observations of transcriptional stimulation of RP genes upon Tail depletion^42^.

We further analyzed changes following coactivator depletion by comparing changes in bulk transcription, F_on_, MPC and Fano at two specific high expression, CR and MED-Tail dependent genes. Depletion of either MED-Tail or SAGA results in large changes in bulk transcription for the *GAS1* gene (**Fig 6C**), and these changes appear to be primarily driven by changes in F_on_. Conversely, at the *AHP1* gene, SAGA depletion results in changes to both F_on_ and MPC but little change in Fano, whereas Tail depletion results in larger decreases in both MPC and Fano with smaller changes to F_on_ (**Fig 6D**). Interestingly, *GAS1* contains a TATA-less promoter while *AHP1* contains a promoter with a TATA box, further strengthening the connection between coactivator complexes, core promoter elements, and transcriptional bursting. Taken together, our combined results demonstrate that MED Tail and SAGA function display a strong connection to F_on_ and a modest connection to MPC and Fano, although their roles in promoting these transcriptional behaviors can vary substantially among affected genes.

## Discussion

Here we present the results of a time-resolved scRNA-seq approach to study transcription parameters in yeast cells. We demonstrate that temporal information gained through this approach is consistent across increasing time courses, and from comparing samples from the same time point across experiments. Consistent with prior work, we found that most genes display relatively low levels of transcriptional variance, while a small subset of genes display exceptionally high variance. We also observe that gene features including core promoter sequence (e.g. TATA boxes), and transcriptional coactivator class correlate with higher Fano values, which agrees well with observations using microscopy data on select model yeast genes^3^. Fano is generally higher for CR genes compared to TFIID genes across expression quartiles and is especially high for cell cycle regulated CR genes.

Current models of gene activation state that generally, the primary mechanism by which transcription is controlled is via modulation of the fraction of active promoters over time. In higher eukaryotes, this has been studied in the context of enhancer-promoter communication, in which an enhancer functions in part to increase the frequency of target promoter activation. An alternative (though not mutually exclusive) form of regulation is modulation of the bursting properties of an already active promoter, regulating the amount of transcription per cell by controlling the duration of the active state and/or the frequency of initiation during the active state. Our results suggest that both of these mechanisms operate for the cell cycle-regulated histone genes that appear to switch transcription from a low level constitutive mode in M and M/G1 to an activated and bursty mode in G1, S, and G2. In our coactivator depletion experiments, we found that the change in fraction of active promoters closely correlates with the bulk transcriptional fold change, consistent with the model where SAGA and Med Tail modulate the fraction of active promoters. We do, however, observe modest changes in mean expression per cell (MPC) (a measure of transcription produced during the active state) after MED Tail or SAGA depletion, suggesting that both Mediator and SAGA can play a modest role in modulation of the bursting properties of an already active promoter. Furthermore, the relative changes in F_on_ vs. MPC can vary widely by gene, suggesting that mechanistically these coactivator complexes likely play different roles in promoting transcription in a gene-dependent fashion. Although we cannot directly address this hypothesis using our dataset, it is conceivable that these coactivators may support more frequent or longer-lived activation periods at some genes, and/or more transcription per active period at others.

## Limitations of the study

Due to limitations in capturing efficiency of RNAs using single cell RNA-seq approaches and metabolic labeling efficiency, our estimates of F_on_ are likely underestimates. In addition, we are not able to distinguish multiple potential activation periods during the metabolic labeling period.

## Supplemental Table and Figure Legends

**Table S1. Estimated transcript half-lives**

**Table S2. Estimated Fano values**

**Table S3. Strains used in this study**

**Figure S1.** Measurement of genome-wide mRNA stability.

**A)** Pearson’s correlation in transcript half-life estimates from different 4-TU labeling times in our dataset (5min, 10min, 15 min), compared to polyA transcript half-lives reported in Chan et al. 2018.^43^ **B)** Correlation between codon adaptation index (CAI) and estimated transcript half lives (10 min timepoint). CAI values reported in Drummund et al.^44^

**Figure S2.** Reproducibility of F_on_ estimates.

**A-F)** Pearson’s correlation comparing F_on_ estimates for labeling time course samples (5 min, 10 min, 15 min) in WT cells **(A-C)** and 10-minute labeling timepoint samples of WT compared with the indicated degron strains without auxin treatment **(D-F)**.

**Figure S3.** New transcript count estimates and total transcript counts per cell

Distributions of new transcript counts and total transcript counts per cell for no 4-TU control cells and 5 min, 10 min, and 15 min 4-TU treated cells. Red dashed lines represent the median transcript count per cell.

**Figure S4.** Range and reproducibility of Fano estimates

**A)** Pearson’s correlation comparing Fano estimates for time course samples (5 min, 10 min, 15 min) and 10 minute labeling timepoint samples without auxin treatment in Med15 and Spt3/7 degron strains. **B)** Fano values (10 min sample) for genes with short (< 3 min) or long (>3 min) TFIIF promoter residence

**Figure S5.** F_on_ for cell cycle regulated and Fano for non-cyclic transcripts

**A)** Log-F_on_ values estimated at each cell cycle stage, comparing groups of transcripts peaking at each cell cycle stage ^40^. **B)** Log Fano values for non-cyclic transcripts, comparing Fano values estimated at each cell cycle stage.

**Figure S6.** Western blot analysis and changes in Fon, MPC and Fano following Med15 or Spt3/7 depletion

**A)** Western blot analysis of coactivator subunits (top) and loading control (bottom) following 30 minute auxin treatment (+) or control DMSO treatment (-). Molecular size standards indicated on left. Upper blot probed with anti V4 (degron tag) or anti Tfg2 (bottom). **B-C)** Log2-fold changes in F_on_, MPC, and Fano for coactivator-regulated gene classes after rapid depletion of Med15 **(B)** or Spt3/7. **(C)**. Gene classes noted are coactivator redundant (CR), MED-Tail dependent (TD) or independent (TI), and TFIID. Statistical test performed is Wilcoxon rank sum test. For all figure panels, significance levels are defined as ∗∗∗∗p ≤ 0.0001, ∗∗∗p ≤ 0.001, ∗∗p ≤ 0.01, ∗p ≤ 0.05, ns p > 0.05.

**Figure S7.** Log2-fold change comparisons of F_on_, MPC, Fano, and bulk transcription following coactivator depletion

**A)** Pairwise comparisons between log2-fold changes in bulk transcription, F_on_, MPC and Fano following acute depletion Med15 (MED Tail) subunit for MED Tail-dependent genes, as in Fig 4. R^2^ values shown are adjusted R-squared values derived from linear model fits. (**B-C)** Pairwise comparisons as in (a) but following acute depletion of SAGA (Spt3/7 subunits) for **(B)** CR genes or **(C)** TFIID genes.

**Figure S8.** Log2-fold changes in bulk transcription, MPC, F_on_ and Fano for ribosomal gene classes following coactivator depletion

**A-B)** Log2-fold change in bulk transcription, MPC, F_on_ and Fano for ribosome biogenesis (RiBi) and ribosomal protein (RP) genes following **(A)** Spt3/7 depletion or **(B)** Med15 depletion. Non-RiBi/RP genes (Other) are shown for comparison. Statistical test performed is Wilcoxon rank sum test. For all figure panels, significance levels are defined as ∗∗∗∗p ≤ 0.0001, ∗∗∗p ≤ 0.001, ∗∗p ≤ 0.01, ∗p ≤ 0.05, ns p > 0.05.

## Author Contributions

**JAS:** conceptualization, methodology, investigation, formal analysis, writing – original draft; review and editing; **SH**: supervision, writing - review & editing; funding acquisition.

## Acknowledgements

We thank Manu Setty, Dan Larson, and members of the Hahn and Tsukiyama labs for their insightful comments on this work. We also thank Fred Hutch Genomics for assisting in sample preparation and consulting on the project development, especially Elizabeth Jensen and Dolores Covarrubias. Supported by NIH grants R35 GM140823 to SH and P30 CA015704 to the Fred Hutch Genomics and Computational Shared Resources facility.

## STAR METHODS

### Time course

WT yeast cells (SHY772) were grown to mid-log phase (OD ∼0.8) and treated with 5 mM 4-TU for 5min, 10min, or 15min or left untreated (DMSO only). Immediately following 4-TU treatment, cells were placed on ice and 2 volumes of ice-cold PBS was added to the culture in 50mL Falcon tubes. Cells were pelleted for 5 min at 3000xg in a benchtop centrifuge at 4degC in a swinging bucket rotor and washed with cold PBS + 0.01% BSA. Cells were pelleted at 3000xg at 4degC for 5min and resuspended in 1ml PBS + 0.01% BSA. Cells were fixed by continuous gentle vortexing in a 50mL Falcon tube while adding ice cold methanol dropwise to a final concentration of 80%. Cells were incubated on ice for 15min, at which point fixed cells were stored overnight at –20degC. The following day, fixed cells were incubated on ice for 15min, then pelleted for 5 min at 3000xg in a benchtop centrifuge at 4degC. Cells were resuspended in 1mL ice cold PBS + 0.01% BSA, transferred to a 1.5mL Eppendorf tube and pelleted at 2000xg for 1min. Cells were resuspended in 400ul PBS + 0.01% BSA, and the cell suspension was added to 1mL of alkylation buffer, followed by a 45min incubation at 45C with occasional inversion. Cells were pelleted at 2000xg for 30sec and resuspended in 1mL PBS + 0.01% BSA + 10mM DTT. This reducing wash step was repeated for a total of two washes, followed by pelleting at 2000xg for 30sec and resuspension of the pellet in 1M sorbitol + 0.01% BSA. Cells were pelleted at 2000xg for 30sec, and cells were then resuspended in 900μl spheroplasting buffer, and 100μl of 100mg/ml Zymolyase 100T was added. Cells were spheroplasted for ∼30min at 30C with occasional inversion. The spheroplasting mixture was filtered through a MACS SmartStrainer and collected in a 1.5mL Eppendorf tube on ice, and residual spheroplasts were rinsed through the SmartStrainer using 500μL ice cold spheroplasting buffer. Spheroplasts were pelleted for 5min at 2000xg forming a cushion of spheroplasts towards the bottom of the Eppendorf tube. Supernatant was removed leaving ∼100ul of spheroplast cushion, which was diluted 1:5 in ice cold 1M sorbitol + 0.01% BSA. Spheroplasts were counted using a hemocytometer immediately prior to Gel Beads in emulsion (GEM) generation.

Coactivator depletion studies were performed in Med15-degron (SHY1055) or Spt3/7-degron (SHY1176) strains. Prior to 4-TU labeling for 10min, cells were treated with either 500μM 3-indoleacetic acid (IAA) in DMSO or DMSO alone as a negative control, and the remainder of the above protocol was performed.

### Western blot analysis

1 ml of each Med15-degron and Spt3/7-degron cell cultures were collected immediately after treatment with IAA or DMSO. Cells were pelleted, then resuspended in 200μl 0.1M NaOH and incubated for 5min at room temperature. Cells were pelleted, then resuspended in 100μl yeast whole cell extract buffer and heated for 5min at 95C. Samples were centrifuged for 5min at max speed, then extracts were subjected to SDS-PAGE using a precast NuPAGE 4%-12% Bis-Tris gel using MOPS running buffer. Proteins were transferred onto a PVDF membrane, and after transfer the membrane was cut at the ladder band (SeeBlue Plus2 pre-stained protein standard) corresponding to 62kDa. A mouse monoclonal primary a-V5 antibody was used to probe Med15-3xV5-IAA7, Spt3-3xV5-IAA7 and Spt7-3xV5-IAA7 fusion proteins, and rabbit polyclonal primary a-Tfg2 antibody was used to probe endogenous Tfg2 as a loading control. Dye-conjugated mouse or rabbit secondary antibodies were used to visualize antibody-bound proteins, and dye signals were captured using an Odyssey CLx scanner.

### Library preparation and sequencing

GEM generation and single cell RNA-seq library preparation was performed using the Chromium Next GEM Single Cell 3’ Kit from 10X Genomics according to manufacturer’s instructions. Prepared libraries were sequenced on an Illumina NextSeq 2000 instrument using P3 reagents, with read configuration of 28nt for R1 to identify cell barcode and UMIs, and 125nt for R2 to sequence the captured mRNA.

### Sequence processing

Single cell IDs were determined from sequencing data using the default CellRanger analysis pipeline (count), using a yeast transcriptome built from 3’-end annotations^45^. Sequencing alignment and T-to-C mutation counting was performed using the TimeLapse-seq computational pipeline (bitbucket link). Custom scripts were used to assign mutation-counted reads with their cells of origin using the cell barcodes in the CellRanger output .bam file. For the time course datasets with higher numbers of input cells, potential doublet cells were identified and filtered using doubletFinder_v3.

### Estimation of transcript half lives

To determine transcript half-lives, we first pooled mutation-called reads from our single cell experiments to create a pseudo-bulk datasets. From these pooled reads, we estimated the fraction of new RNAs per gene over each labeling period using a Bayesian modeling approach (bakR^35^). bakR was used to estimate per-sample and background (from no 4TU-treated control) T-to-C mutation rates, then was used to fit per-gene fraction new for each timepoint using an maximum likelihood estimation (fast_analysis). We then calculated transcript half-lives using the following equation:

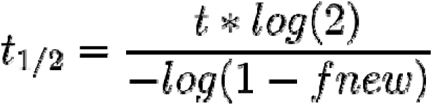

where t is the labeling time and fnew is the estimated fraction new.

### Estimation of per gene per cell new RNA counts

To account for potential background mutations in sequencing, nascent read counts were simulated from the subset of T-to-C mutation containing reads using a binomial probability approach. Each read was assigned a probability of being newly transcribed based on the global sample mutation rate, number of Ts in the read, and number of T-to-C mutations in the read. Each read was then subjected to a random draw according to its binomial probability to assign it as new or pre-existing. Simulated new reads were then summed per-gene and per-cell to generate the final new read count matrix.

### Fano calculation

For each gene, the mean (*m*) and variance (σ) of estimated new read counts across cells were calculated. Fano was then calculated according to the following equation: Fano = σ^2^/*m*.

### Estimation of per gene fraction on (F_on_)

To estimate the fraction of cells active over the labeling period for each gene, we estimated gene-specific mutation rates for all reads in each single cell by summing the total number of T-to-C mutations per gene per cell, and dividing by the total number of Ts observed per gene per cell. We then assigned a cell as actively expressing a given gene over the label period if this mutation rate exceeded ten-fold above the background mutation rate derived from a non-labeled control. F_on_ was then calculated by dividing the number of active cells per gene by the total number of cells.

### Cell cycle assignments

Cells were assigned to a cell cycle state through k-means clustering of new read counts of highly cell cycle regulated genes. Briefly, a matrix of new RNA counts for genes vs. cells was filtered to contain only cell cycle regulated genes with a high periodicity score determined in Spellman et al.^40^ (n = 297). K-means clustering was performed to group cells into five clusters, corresponding to G1, S, G2, M, and M/G1 cell cycle stages.

### Buffers

Alkylation buffer (1ml) (700μl DMSO, 140μl 500mM sodium phosphate pH 8, 140μl 100mM iodoacetamide, 20μl water) Spheroplasting buffer (1ml) (500μl 2M sorbitol, 350μl water, 100μl 1% BME, 20μl 500mM EDTA pH 8, 20μl 500mM sodium phosphate pH 8, 10μl 10mg/ml BSA) Yeast whole cell extract buffer (0.06M Tris-HCl, pH 6.8, 10% glycerol, 2% SDS, 5% 2-mercaptoethanol, 0.0025% bromophenol blue)

## Data Availability

High-throughput sequencing data will be available after peer review.

## Code Availability

Scripts used to process datasets in this manuscript are available by request from the lead contact without restriction.

