## supplemental figures S1-S8 for "Transcriptional bursting, gene activation, and roles of SAGA and Mediator Tail measured using nucleotide recoding single cell RNA-seq"

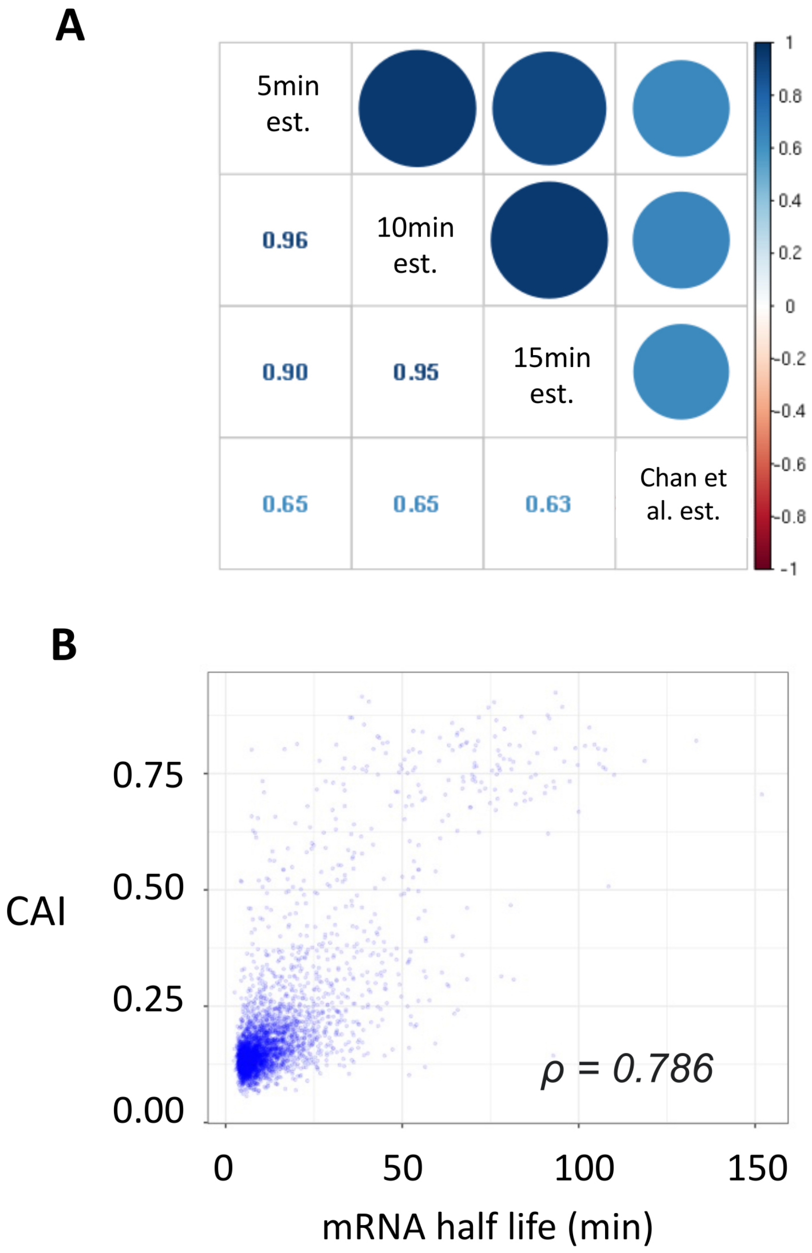


**Figure S1.** Measurement of genome-wide mRNA stability.
**A)** Pearson's correlation in transcript half-life estimates from different 4-TU labeling times in our dataset (5min, 10min, 15 min), compared to polyA transcript half-lives reported in Chan et al. 2018.[43] **B)** Correlation between codon adaptation index (CAI) and estimated transcript half-lives (10 min timepoint). CAI values reported in Drummund et al.[44]

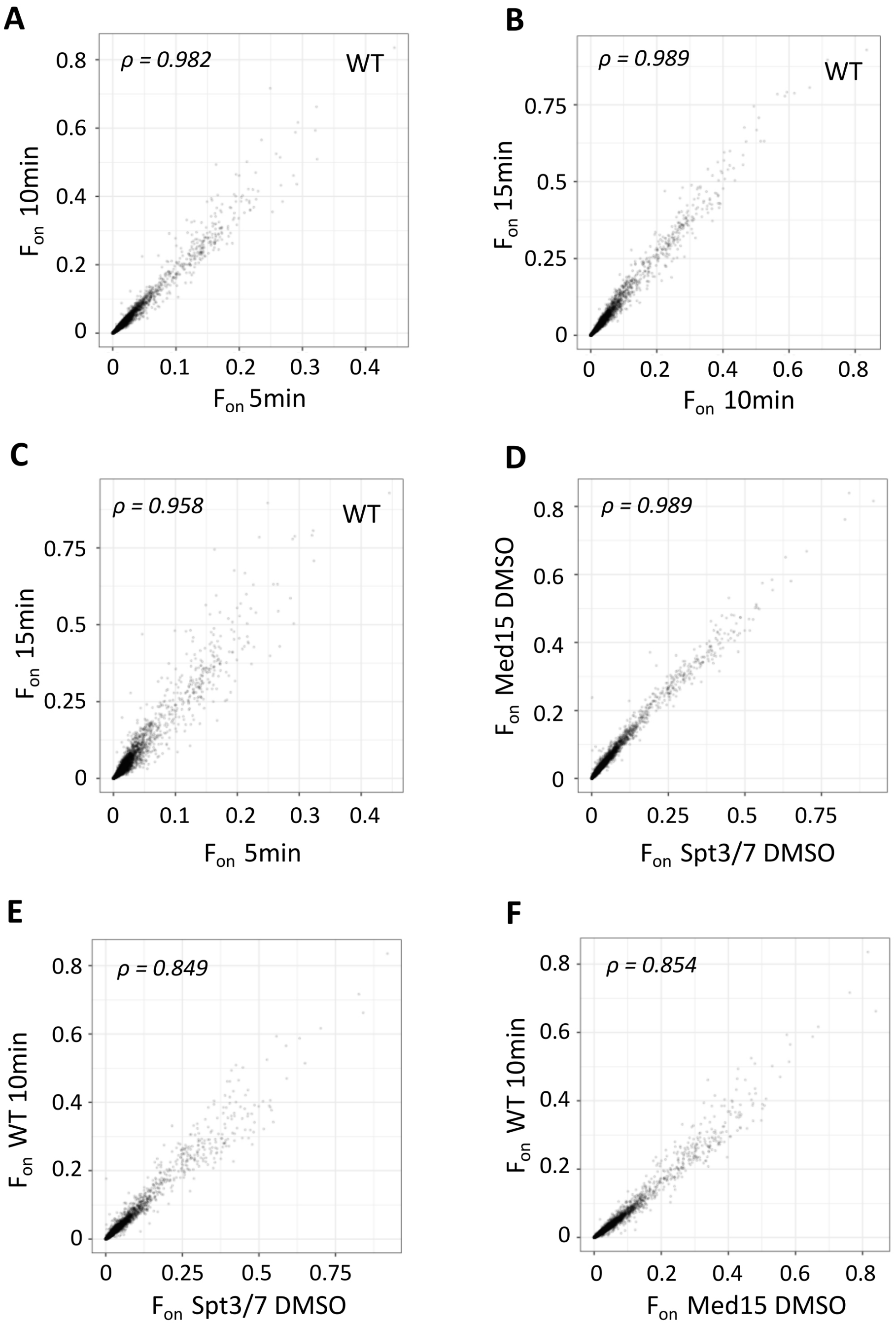


**Figure S2.** Reproducibility of $F_{on}$ estimates.

**A-F** Pearson's correlation comparing $F_{on}$ estimates for labeling time course samples (5 min, 10 min, 15 min) in WT cells **(A-C)** and 10-minute labeling timepoint samples of WT compared with the indicated degron strains without auxin treatment **(D-F)**.

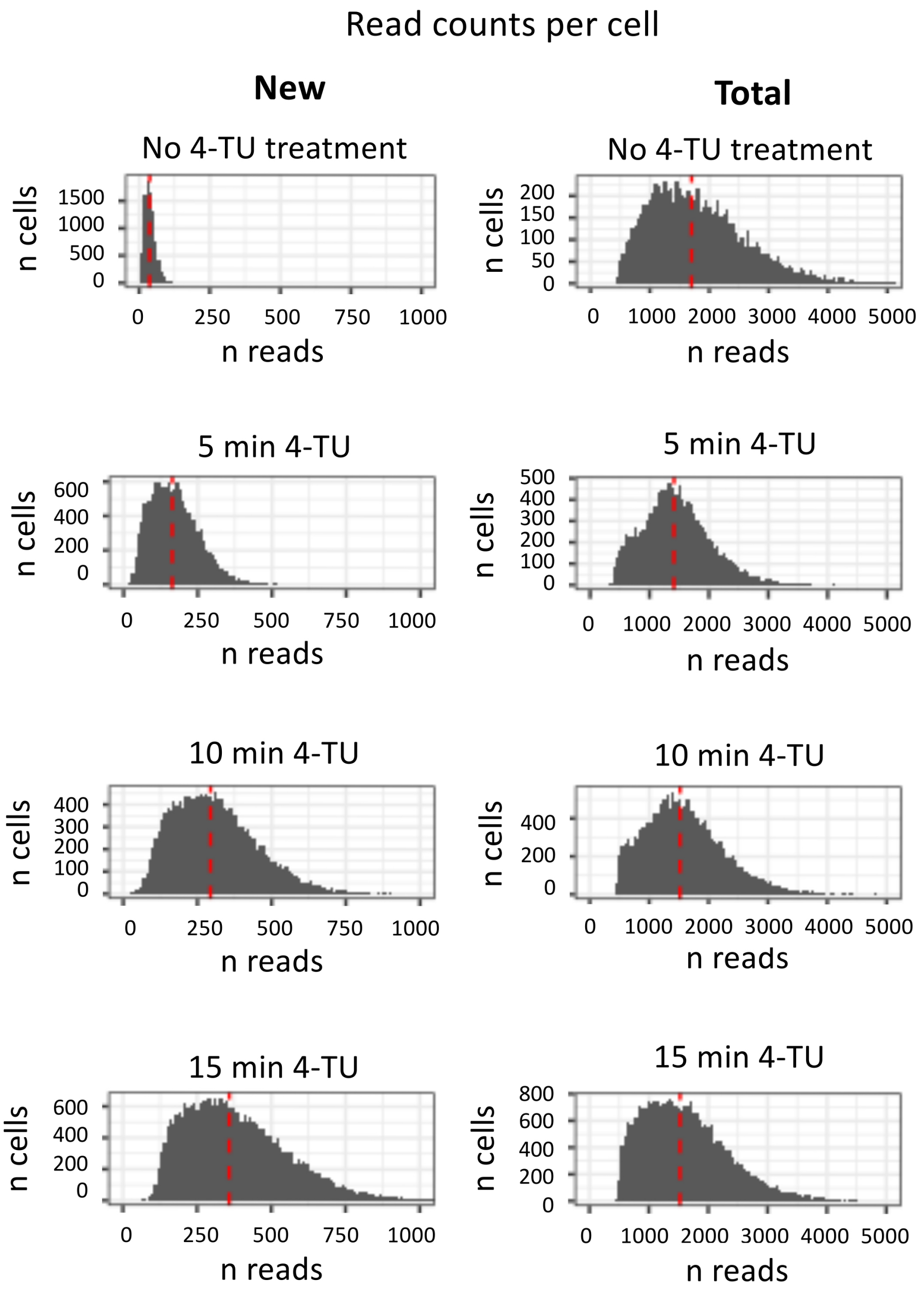


**Figure S3.** New transcript count estimates and total transcript counts per cell
Distributions of new transcript counts and total transcript counts per cell for no 4-TU control cells and 5 min, 10 min, and 15 min 4-TU treated cells. Red dashed lines represent the median transcript count per cell.

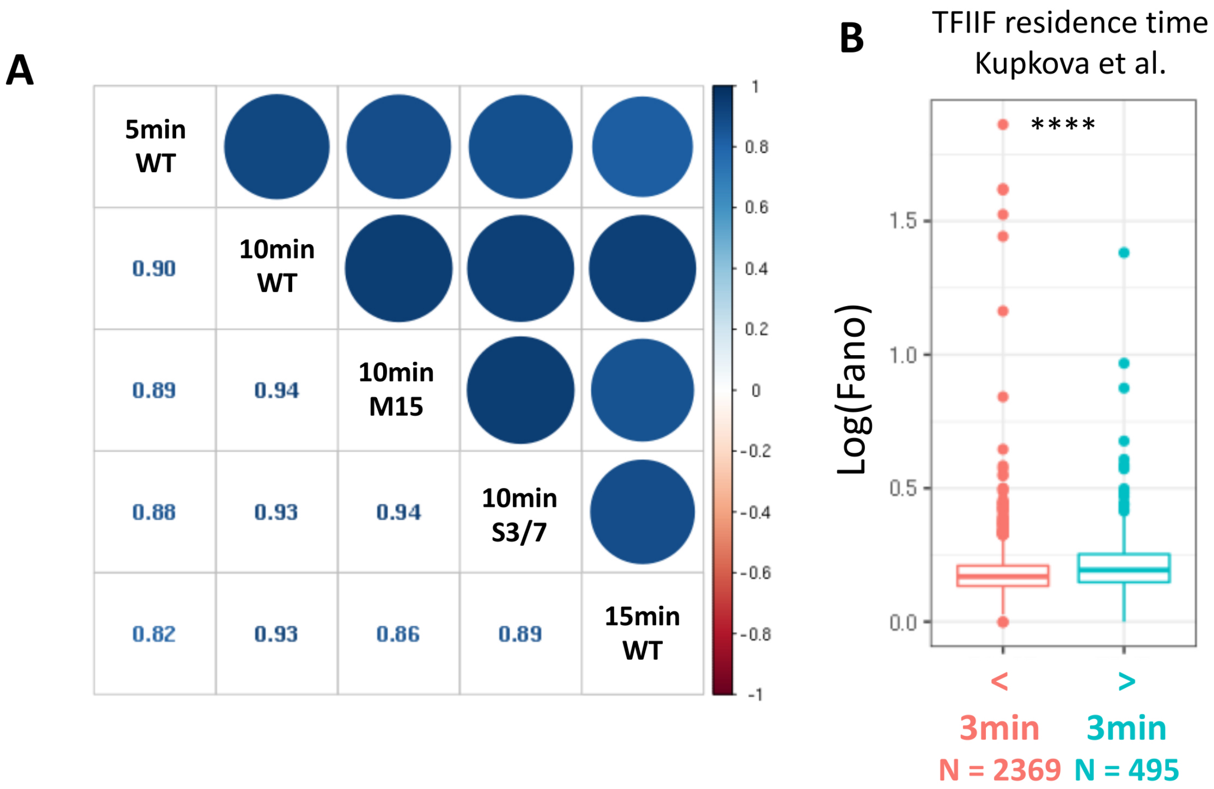


**Figure S4.** Range and reproducibility of Fano estimates
**A)** Pearson's correlation comparing Fano estimates for time course samples (5 min, 10 min, 15 min) and 10 minute labeling timepoint samples without auxin treatment in Med15 and Spt3/7 degron strains. **B)** Fano values (10 min sample) for genes with short (< 3 min) or long (>3 min) TFIIF promoter residence time.[39]

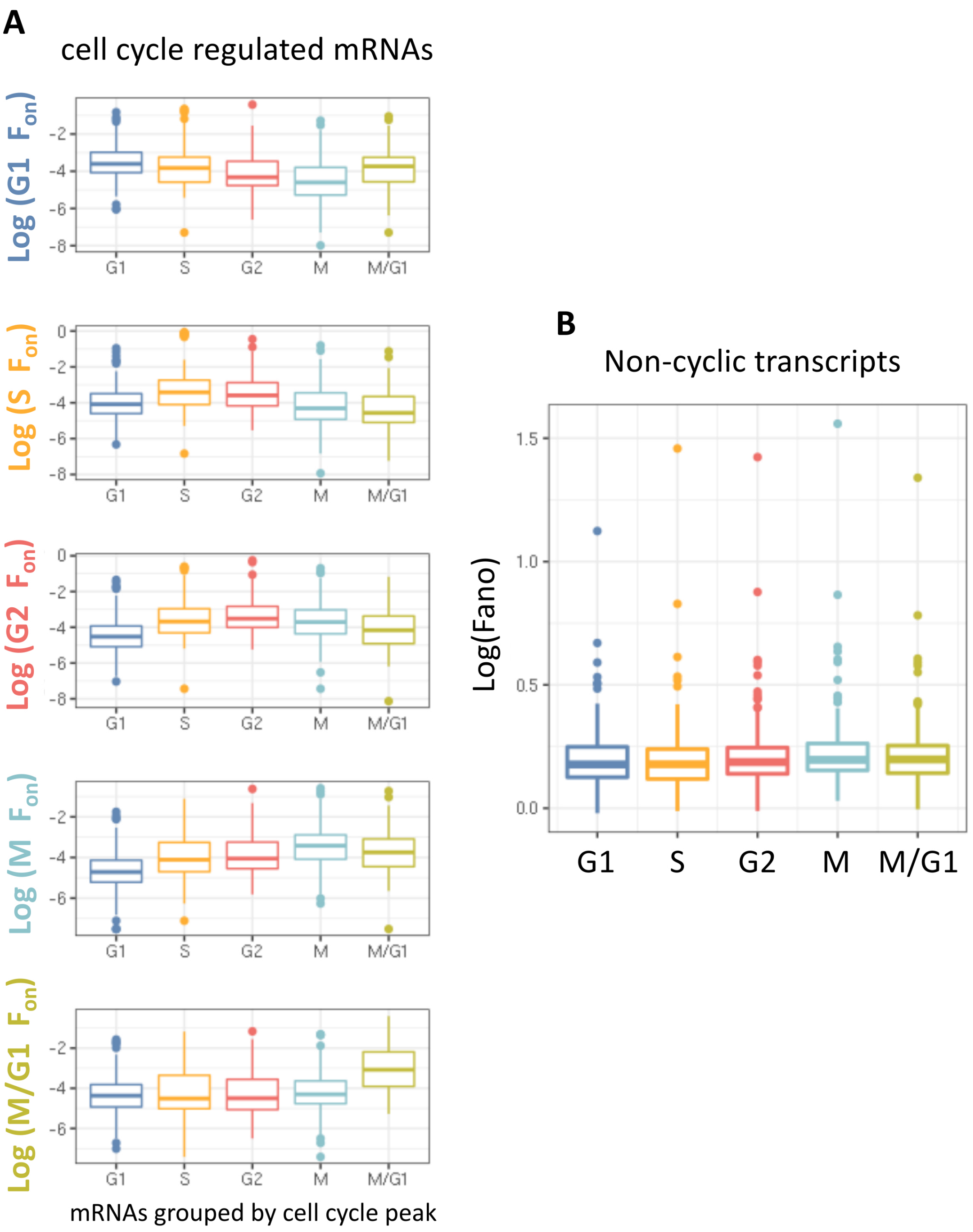


**Figure S5**. $F_{on}$ values for cell cycle regulated transcripts and Fano values for non-cyclic transcripts
**A)** Log-$F_{on}$ values estimated at each cell cycle stage, comparing groups of transcripts peaking at each cell cycle stage [40]. **B)** Log Fano values for non-cyclic transcripts, comparing Fano values estimated at each cell cycle stage.

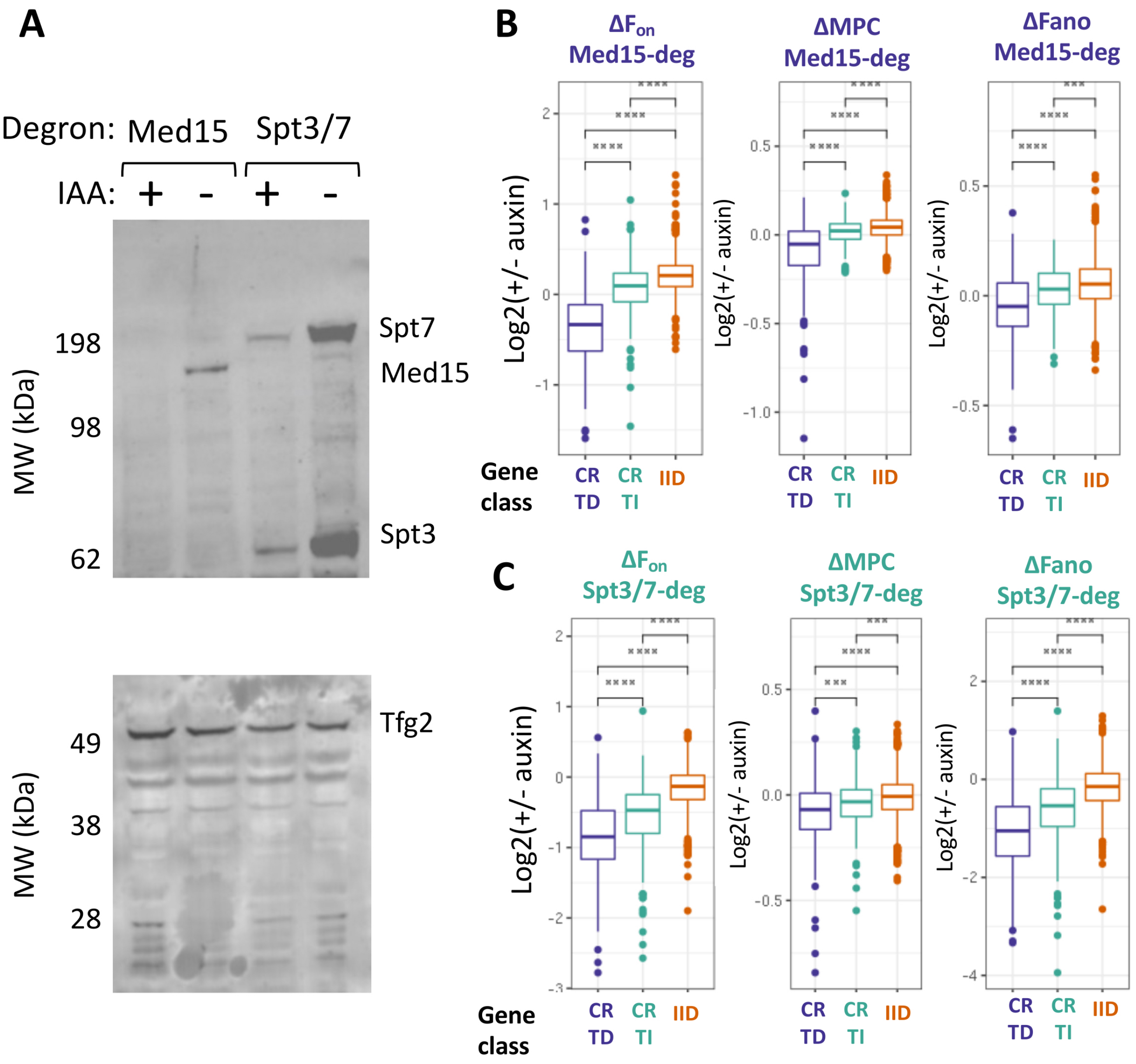


**Figure S6.** Western blot analysis and changes in Fon, MPC and Fano following Med15 or Spt3/7 depletion

**A)** Western blot analysis of coactivator subunits (top) and loading control (bottom) following 30 minute auxin treatment (+) or control DMSO treatment (-). Molecular size standards indicated on left. Upper blot probed with anti V4 (degron tag) or anti Tfg2 (bottom). **B-C)** Log2-fold changes in $F_{on}$, MPC, and Fano for coactivator-regulated gene classes after rapid depletion of Med15 **(B)** or Spt3/7. **(C)**. Gene classes noted are coactivator redundant (CR), MED-Tail dependent (TD) or independent (TI), and TFIID. Statistical test performed is Wilcoxon rank sum test. For all figure panels, significance levels are defined as ****p ≤ 0.0001, ***p ≤ 0.001, **p ≤ 0.01, *p ≤ 0.05, ns p > 0.05.

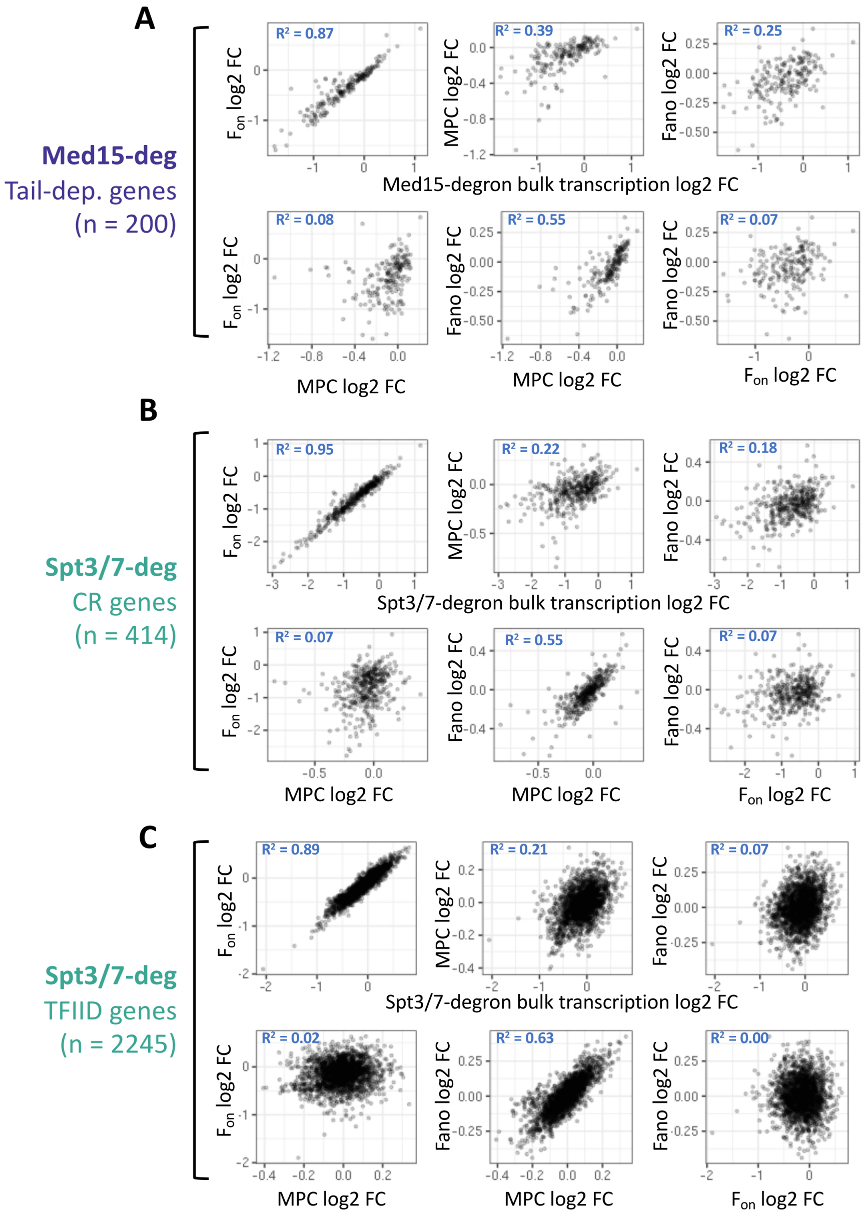


**Figure S7.** Log2-fold change comparisons of $F_{on}$, MPC, Fano, and bulk transcription following coactivator depletion

- **A)** Pairwise comparisons between log2-fold changes in bulk transcription, $F_{on}$, MPC and Fano following acute depletion Med15 (MED Tail) subunit for MED Tail-dependent genes, as in Fig 4. $R^2$ values shown are adjusted R-squared values derived from linear model fits. (**B-C)** Pairwise comparisons as in (a) but following acute depletion of SAGA (Spt3/7 subunits) for **(B)** CR genes or **(C)** TFIID genes.

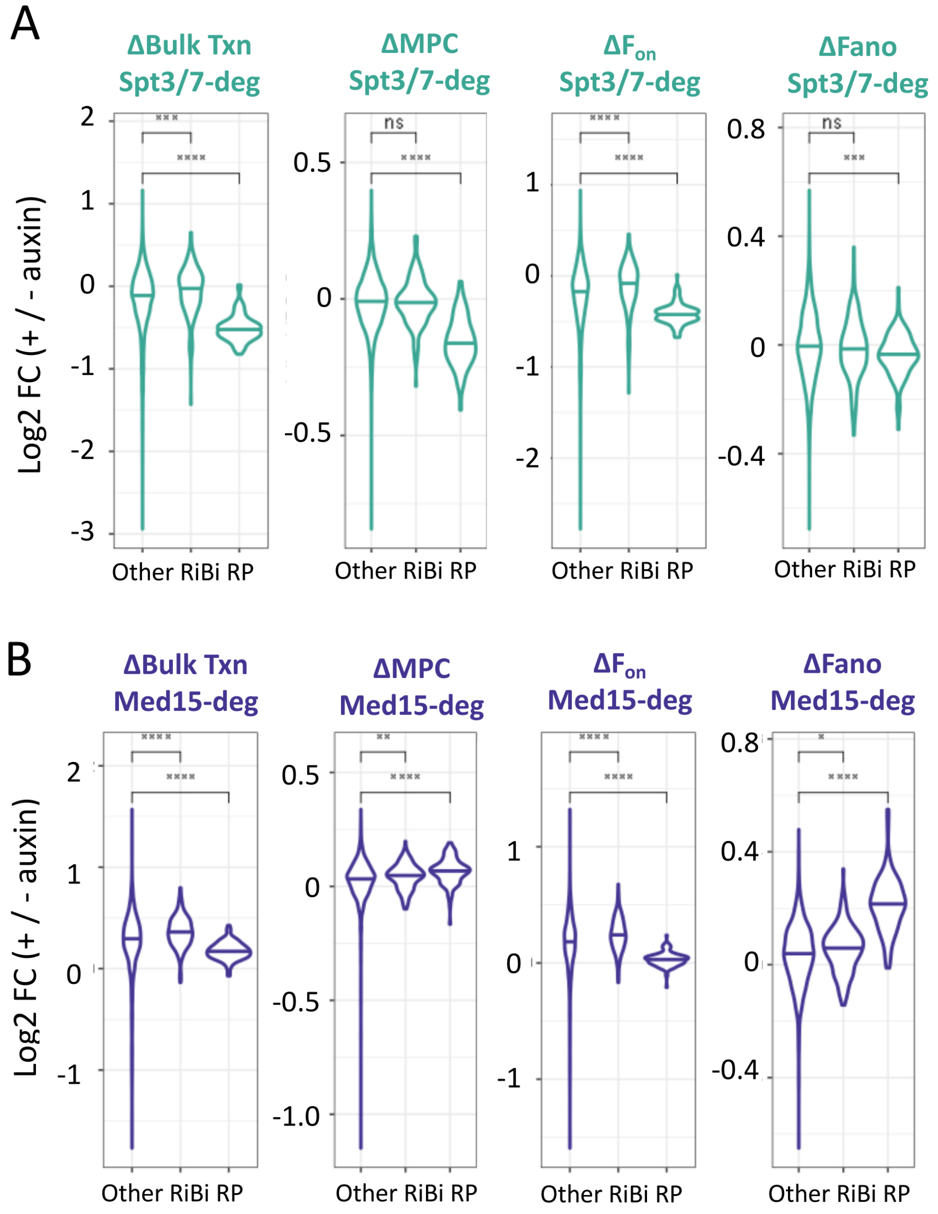


**Figure S8**. Log2-fold changes in bulk transcription, MPC, $F_{on}$ and Fano for ribosomal gene classes following coactivator depletion

**A-B)** Log2-fold change in bulk transcription, MPC, $F_{on}$ and Fano for ribosome biogenesis (RiBi) and ribosomal protein (RP) genes following **(A)** Spt3/7 depletion or **(B)** Med15 depletion. Non-RiBi/RP genes (Other) are shown for comparison. Statistical test performed is Wilcoxon rank sum test. For all figure panels, significance levels are defined as $****p \leq 0.0001$, $***p \leq 0.001$, $**p \leq 0.01$, $*p \leq 0.05$, ns $p > 0.05$.
